# Protein homeostasis from diffusion-dependent control of protein synthesis and degradation

**DOI:** 10.1101/2023.04.24.538146

**Authors:** Yuping Chen, Jo-Hsi Huang, Connie Phong, James E. Ferrell

## Abstract

It has been proposed that the concentration of proteins in the cytoplasm maximizes the speed of important biochemical reactions. Here we have used the *Xenopus* extract system, which can be diluted or concentrated to yield a range of cytoplasmic protein concentrations, to test the effect of cytoplasmic concentration on mRNA translation and protein degradation. We found that protein synthesis rates are maximal in ∼1x cytoplasm, whereas protein degradation continues to rise to an optimal concentration of ∼1.8x. This can be attributed to the greater sensitivity of translation to cytoplasmic viscosity, perhaps because it involves unusually large macromolecular complexes like polyribosomes. The different concentration optima sets up a negative feedback homeostatic system, where increasing the cytoplasmic protein concentration above the 1x physiological level increases the viscosity of the cytoplasm, which selectively inhibits translation and drives the system back toward the 1x set point.

## Introduction

The cytoplasm is crowded with macromolecules, with proteins being the most abundant class. In total, proteins constitute about 82% of the dry mass of a mammalian cell (Oh et al., 2022) and about 55% of an *E*. *coli* cell (Milo et al., 2009), and the cytoplasmic concentration of protein ranges from about ∼75 mg/mL in mammalian cell lines (Liu et al., 2022) and *Xenopus* egg extracts (Green, 2009; Murray, 1991) to 200-320 mg/mL in *E*. *coli* (Elowitz et al., 1999; Zimmerman and Trach, 1991). Assuming an average protein density of 1.35 g/mL, this means that the volume fraction of a cell occupied by protein is ∼6% to 25%, and the macromolecular excluded volume is higher still. Although protein concentration transiently decreases during mitosis (Son et al., 2015; Zlotek-Zlotkiewicz et al., 2015), for a given cell type, the concentration of macromolecules in the cytoplasm is tightly regulated and remarkably constant (Delarue et al., 2018; Liu et al., 2022; Neurohr and Amon, 2020; Oh et al., 2022).

This brings up two basic questions: why is the cytoplasmic protein concentration as high as it is, and no higher, and what mechanisms set and maintain this concentration? It has been conjectured that the normal cellular protein concentration maximizes the rates of important biochemical reactions, with there being a trade-off between the effects of concentration on enzyme-substrate proximity and viscosity (Dill et al., 2011). This conjecture assumes that these critical reactions are sensitive to the viscosity of the medium because the collision frequency of their reactants depends upon viscosity. Note, however, that decades of in vitro work has indicated that almost all enzymes are reaction-limited, not diffusion-limited (Bar-Even et al., 2011), so that changes in diffusion might be expected to have minimal effect on the rates of most enzyme reactions. That said, there are examples of protein-mediated processes that are markedly inhibited by increased viscosity (Alric et al., 2022; Drenckhahn and Pollard, 1986; Molines et al., 2022; Tan et al., 2013), and the simplest explanation for this is that they are diffusion controlled. Thus, the maximal speed conjecture remains an intriguing possibility.

Here we set out to directly test the conjecture experimentally. We chose to use the *Xenopus* egg extract system, an undiluted cell-free living cytoplasm, for these studies because they allow easy, direct manipulation of the concentration of cytoplasmic macromolecules. Although *Xenopus* extracts are an in vitro system, they carry out the complex biological functions of an intact *Xenopus* egg or embryo remarkably faithfully. For example, extracts can self-organize into sheets of cell-like structures whose overall architecture closely resembles that of embryonic blastomeres (Cheng and Ferrell, 2019; Gires et al., 2023; Mitchison and Field, 2021). If supplied with sperm chromatin, extracts can organize a functional nucleus and carry out DNA replication. Indeed, extracts can perform full cell cycles, complete with replication, mitosis, and reductive division of the cell-like cytoplasmic structures (Afanzar et al., 2020; Cheng and Ferrell, 2019; Fang and Newport, 1991; Murray and Kirschner, 1989) .

The processes we chose to examine were protein synthesis and degradation, which are not only potentially affected by the cytoplasmic protein concentration, but also directly involved in determining the protein concentration. *Xenopus* extracts can carry out protein synthesis from their own stores of mRNAs or from added synthetic mRNAs (Kanki and Newport, 1991; Minshull et al., 1989; Ruderman et al., 1979), and they degrade both endogenous proteins (most notably the various substrates of the APC/C) (Kamenz et al., 2021; King et al., 1996) and added probe proteins [the present work]. In some ways, both *Xenopus* extracts and the eggs from which they are derived represent unusual systems for studies of protein control. Transcription is largely absent from fertilized eggs and developing embryos until the midblastula transition (Amodeo et al., 2015; Newport and Kirschner, 1982a, b), and the mRNA content is unchanging throughout the cleavage stages of embryogenesis. In addition, although translational control is critical during oocyte maturation and immediately after fertilization (Fox et al., 1989; Meneau et al., 2020; Paris et al., 1988; Richter et al., 2007; Rosenthal et al., 1983), it appears not to be important during much of embryogenesis (Peshkin et al., 2015). In other ways, the dynamics of proteins in *Xenopus* extracts are typical of animal cells. Common mammalian cell lines have protein concentrations of about 75 mg/mL (Liu et al., 2022), as do *Xenopus* extracts (Murray, 1991), and the median protein half-lives in *Xenopus* extracts (Peshkin et al., 2015) and mouse NIH3T3 cells (Schwanhausser et al., 2011) are almost identical. The simplicity, manipulability, and verisimilitude of the extract system makes it an attractive choice for the present studies of how protein synthesis and degradation are affected by the cytoplasmic concentration.

In this study, we directly manipulated the cytoplasmic macromolecule concentration and viscosity of *Xenopus* egg extracts and assessed the immediate effects on protein synthesis and degradation. We found that translation rates were maximal in 1x cytoplasmic extracts; translation initially increased with cytoplasmic concentration, but then sharply decreased above 1x. Thus, translation is an example of a process whose speed is in fact maximized at the normal concentration of cytoplasmic macromolecules. In contrast, degradation rises with cytoplasmic concentration until the concentration reaches almost 2x. The different concentration optima can be explained by the sensitivities of the processes to viscosity; the viscogen Ficoll 70 was found to be a more potent inhibitor of protein synthesis than of protein degradation. The differential sensitivity of protein synthesis and protein degradation to viscosity provides a simple mechanism for ensuring long-term protein concentration homeostasis.

## Results

### *Xenopus* egg extracts are robust towards cytoplasmic dilution and concentration

The most commonly used types of extract are CSF (cytostatic factor)-arrested extracts, interphase-arrested extracts, and cycling extracts (Cheng and Ferrell, 2019; Dunphy et al., 1988; Murray, 1991; Newport and Spann, 1987). In pilot experiments we found that cycling extracts were the most reliable and longest lived, and so they were chosen for most of the experiments that follow. One caveat is that previous work has shown that translation rates vary between interphase and M-phase (Kanki and Newport, 1991), raising the concern that the time course of protein synthesis would be characterized by alternations between different rates. This proved not to be the case, possibly because mitosis sweeps through the extract in spatial waves (Chang and Ferrell, 2013) (see Movies S1 and S2), so that both mitotic and interphase rates would probably contribute to the measured rate of synthesis at all time points.

To alter the macromolecular concentration of cycling extracts, we used a spin-column with a 10 kilodalton (kDa) cutoff to produce a cytosolic filtrate and a cytoplasmic concentrate (Figure 1B). The filtrate was found to be essentially protein-free by Bradford assay as well as by gel electrophoresis and Trihalo compound or Coomassie staining (Figure 1C and S1). The retentate was on average 2-fold concentrated compared to the starting cytoplasm by Bradford assay (Figure 1C). We then diluted either the starting 1x extract or the 2x retentate with the filtrate to generate extracts with a range of cytoplasmic macromolecule concentrations (Figure 1D, E).

**Figure 1.**
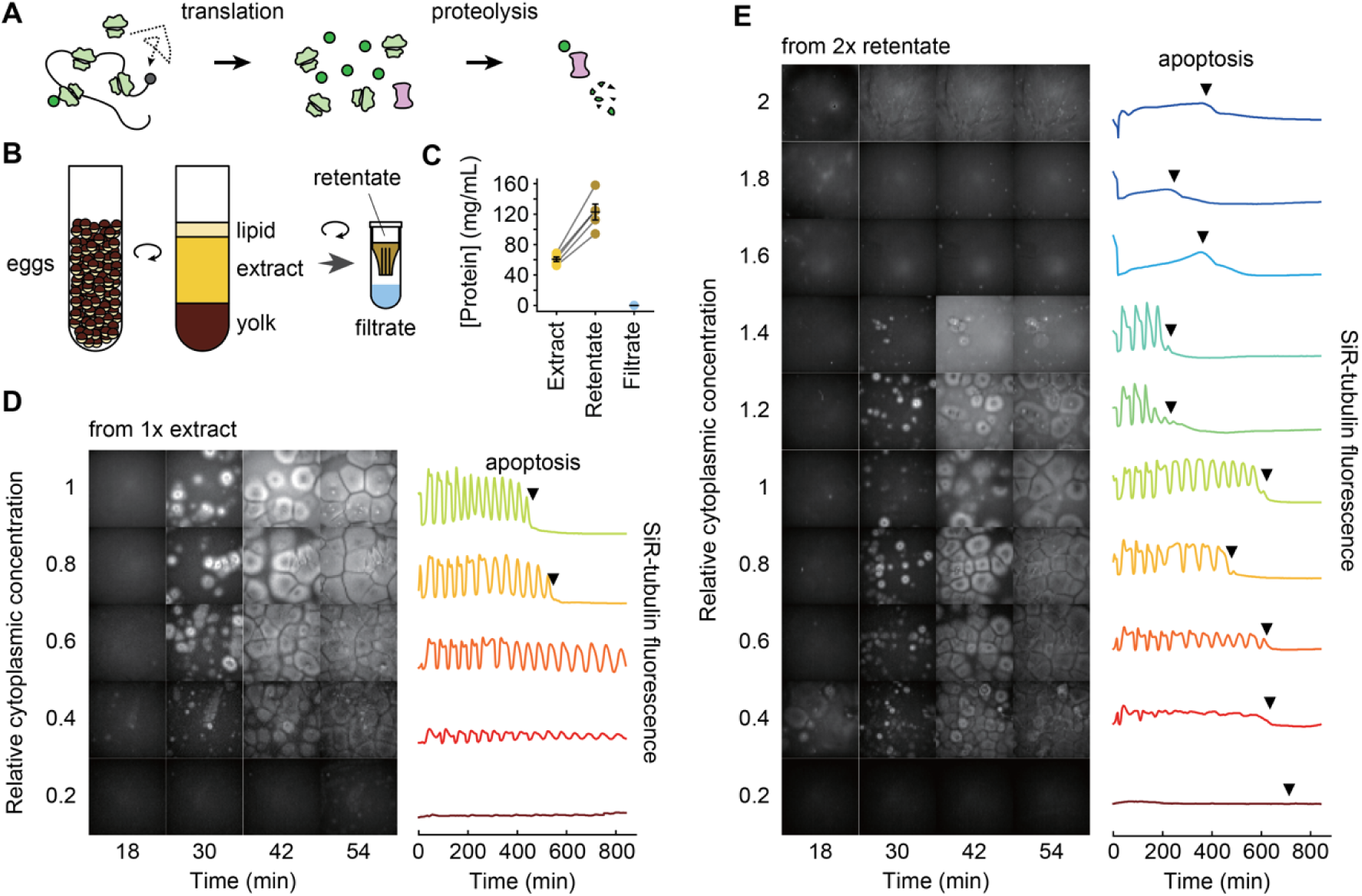
General properties of diluted and concentrated *Xenopus* egg extracts: effects on self-organization and cycling. (A) Schematic view of protein synthesis and degradation. (B) Preparation of *Xenopus* egg extract, 2x concentrated retentate, and protein-depleted filtrate. (C) Protein concentration in extract, retentate, and filtrate. Concentrations were determined by Bradford assays. Data are from five extracts. Individual data points are overlaid with the means and standard errors. (D) SiR-tubulin staining (left) and SiR-tubulin fluorescence intensity as a function of time (right) in an extract after various dilutions. The starting material was a 1x extract, diluted with various proportions of filtrate and imaged in a 96-well plate under mineral oil. All fields are shown at equal exposure. The fluorescence intensities shown on the right were quantified from the center 1/9 of the wells. (E) SiR-tubulin staining (left) and SiR-tubulin fluorescence intensity as a function of time (right) in an extract after various dilutions. The data were collected as in (D) except that the starting material was a 2x extract.

We first examined the effects of concentration on two basic aspects of the extract’s function: its ability to self-organize and to cycle. The extract’s gross organization was found to be remarkably robust to cytoplasmic dilution (Figure 1D, E). Using a microtubule stain, SiR-tubulin, cell-like compartments (Afanzar et al., 2020; Cheng and Ferrell, 2019) were found to form even when the extract was diluted to as low as 0.3x (Figure 1D, E, and Movies S1-S3), in general agreement with a previous report (Cheng and Ferrell, 2019), and to partially organize, with small asters, even at 0.2x the normal cytoplasmic concentration (Movie S3). By following microtubule polymerization and depolymerization, cell cycles were detected down to dilutions of 0.2x, and the oscillations persisted for at least 14 hours with more than 10 complete periods (Figure 1C, D, Movies S1-S3). In agreement with a previous study (Jin et al., 2022), the diluted extracts did not show signs of significant cell cycle defects except that the duration of interphase and the cell cycle period increased with increasing dilution.

On the other hand, the 2x retentate was stickier and more viscous than a 1x extract. Cell-like compartments failed to form at concentrations higher than 1.4x (Figure 1E), and the cell cycle appeared to be arrested above 1.4x. Nevertheless, when a 2x extract was diluted to 1.4x or less, cell-like compartment formation was restored, and the cell cycle behavior was almost normal (Figure 1E), although dilutions from the 2x retentate had slightly longer interphases and were slightly more susceptible to apoptosis than dilutions from a 1x extract (Figures 1D and E).

Overall, extracts were more sensitive to being concentrated than diluted, but even so, extracts were able to carry out grossly normal self-organization and cycling over a wide range of cytoplasmic concentrations.

### Protein synthesis peaks at a 1x cytoplasmic concentration

To measure the protein synthesis rate, we added an mRNA for eGFP and monitored fluorescence as a function of time. In pilot experiments, we titrated the mRNA concentration and found that we could obtain a satisfactory signal without saturating the translation machinery using a concentration of 2.5 ng/µL (Figure 2A). We then recorded time courses of eGFP fluorescence intensity for diluted and concentrated extracts all with the same 2.5 ng/μL concentration of mRNA for eGFP, and calculated translation rates from the linear portion of the time course (Figure 2B). Figure 2C summarizes the data as directly obtained, with equal concentrations of mRNA at each dilution but differing concentrations of the translation machinery. In principle, calculated rates can be plotted against either relative cytoplasmic concentration using the average fold relation from multiple experiments (Figure 1C) or absolute protein concentration measured for each experiment. However, we observed that absolute protein concentration for individual experiments was no better at addressing the variability among experiments than relative cytoplasmic concentration (Figure S2). Furthermore, relative cytoplasmic concentration provides a more straightforward theoretical treatment, as discussed later. Therefore, we opted to use relative cytoplasmic concentration in instances where both relative cytoplasmic concentration and protein concentration were viable options. Translation increased with cytoplasmic concentration to a maximum at ∼0.75x and fell thereafter. To relate this to endogenous translation, given that the endogenous mRNAs would vary with concentration just as the translation machinery does, we calculated an inferred endogenous translation rate, taken as the observed translation rate multiplied by the cytoplasmic concentration. This is shown in Figure 2D; maximal translation was obtained at a cytoplasmic concentration of ∼1x. We also calculated the translation rate normalized for both equal ribosome concentration and equal mRNA concentration, by dividing the raw translation rates by the cytoplasmic concentration (Figure 2E). This provides an estimate of how the specific activity of the translation machinery is affected by cytoplasmic concentration. The inferred specific activity fell markedly with increasing cytoplasmic concentration above 0.5x; below that there was too much variability to draw conclusions. These findings show that protein synthesis is fastest in 1x cytoplasm, as predicted by the maximal speed conjecture (Dill et al., 2011), and suggest that protein synthesis is inhibited when the cytoplasmic viscosity is higher than normal. This latter point is explored further below.

**Figure 2.**
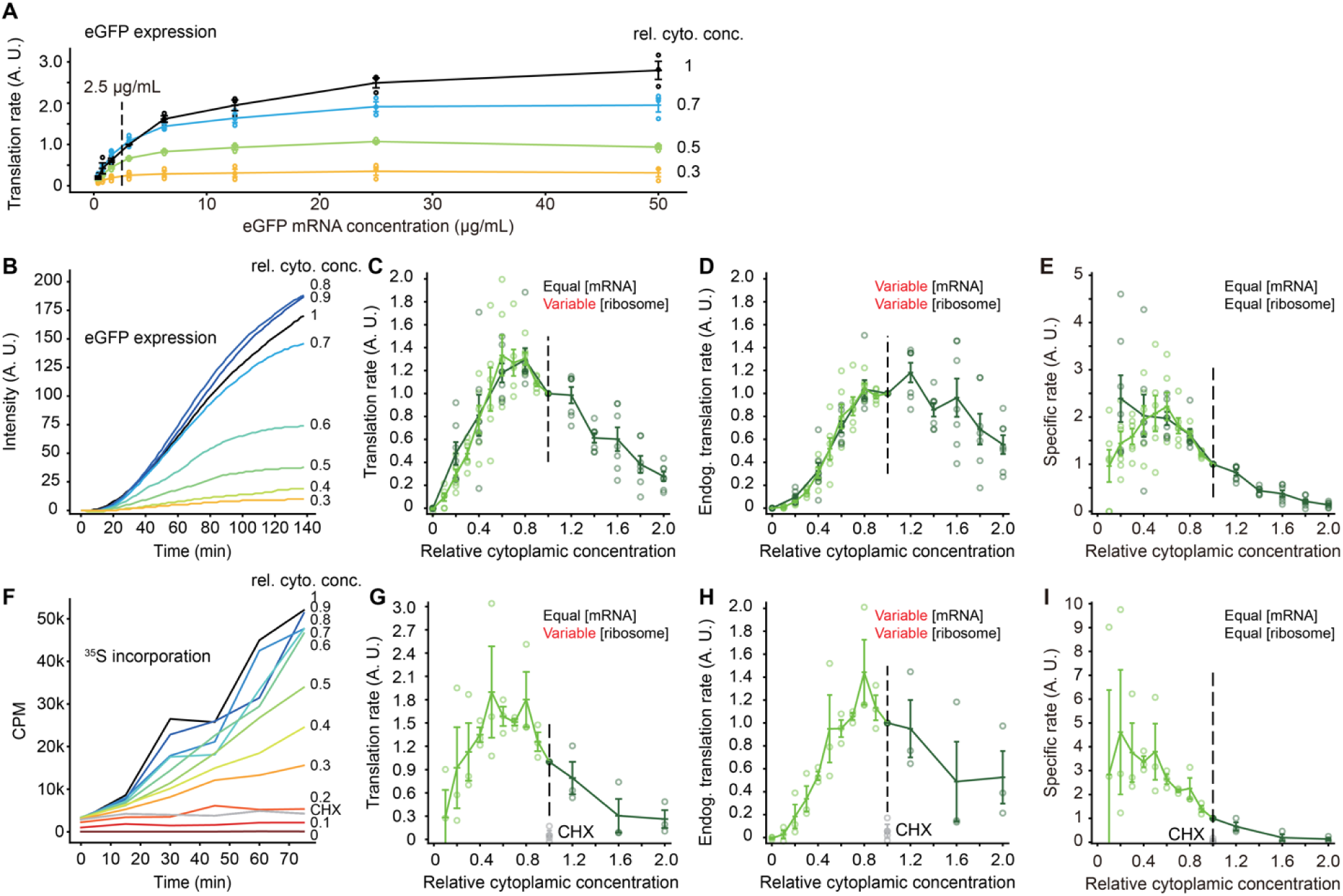
The rate of mRNA translation is maximal at a cytoplasmic concentration of ∼1x. (A) Titration of mRNA concentration for eGFP expression. The indicated concentration (2.5 µg/mL) was chosen for the experiments in (B-E). (B) eGFP expression as a function of time for various dilutions of a 1x extract. (C) Translation rate as a function of cytoplasmic concentration. These are the directly-measured data from experiments where the eGFP mRNA concentration was kept constant and the translation machinery was proportional to the cytoplasmic concentration. Data are from 6 experiments for dilution from 1x extracts and 7 experiments for dilution from 2x retentates. Data are normalized relative to the translation rates at a cytoplasmic concentration of 1x. Means and standard errors are overlaid on the individual data points. In this and the subsequent panels, the darker green represents data from diluting 2x retentates and the lighter green from diluting 1x extract. (D) Inferred translation rates for the situation where the mRNA concentration as well as the ribosome concentration is proportional to the cytoplasmic concentration. The rates from (C) were multiplied by the relative cytoplasmic concentrations. (E) Inferred translation rates for the situation where both the mRNA concentration and the ribosome concentration are kept constant at all dilutions. The rates from (C) were divided by the relative cytoplasmic concentrations. (F) TCA-precipitable ^35^S incorporation as a function of time for translation from endogenous mRNAs. Various dilutions of a 1x extract are shown. CHX denotes a 1x extract treated with 100 µg/mL cycloheximide. (G) Inferred translation rates for the situation where mRNA concentration is kept constant and ribosome concentration is proportional to the cytoplasmic concentration. The rates from (H) were divided by the relative cytoplasmic concentration. The grey data points are from CHX (100 µg/mL)-treated 1x extracts. (H) Translation rate as a function of cytoplasmic concentration. These are the directly-measured data from experiments where the ^35^S concentration was kept constant but both the (endogenous) mRNA concentration and translational machinery were proportional to the cytoplasmic concentration. Data are from 3 experiments for dilution from 1x extracts and 3 experiments for dilution from 2x retentates. Data are normalized relative to the translation rates at a cytoplasmic concentration of 1x. Means and standard errors are overlaid on the individual data points. (I) Inferred translation rates for the situation where both the mRNA concentration and the ribosome concentration are kept constant at all dilutions. The rates from (H) were divided twice by the relative cytoplasmic concentrations (i.e. by the relative concentration squared).

As a second way of gauging the translation rate, we added equal concentrations of ^35^S- methionine to extracts with a range of cytoplasmic concentrations and no added exogenous mRNA, and measured ^35^S incorporation into TCA-precipitable material. Figure 2F shows that incorporation increased linearly with time through 75 min, and that the protein synthesis inhibitor cycloheximide blocked this incorporation. Translation peaked at a cytoplasmic concentration of ∼0.8x (Figure 2G). Overall the dependence of translation on cytoplasmic concentration was very similar to that seen with eGFP (Figure 2D and 2G), and again the inferred specific activity of the translation machinery fell with cytoplasmic concentrations above ∼0.5x (Figure 2H). Thus the effects of cytoplasmic concentration seen with eGFP translation appear to be applicable to translation from endogenous mRNAs as well.

### Protein degradation peaks at a higher cytoplasmic concentration

As a first measure of protein degradation, we made use of an exogenous protein substrate, a heavily BODIPY (boron-dipyrromethene)-labeled BSA, DQ-BSA (for dye-quenched bovine serum albumin). This protein becomes fluorescent during degradation because the BODIPY groups become dequenched. We carried out titration experiments, which showed that an approximately linear response could be obtained with a DQ-BSA concentration of 5 µg/mL (Figure 3A, B). This concentration was then used for experiments with extracts diluted to various extents. As shown in Figure 3C, the rate of DQ-BSA dequenching increased with the concentration of macromolecules and peaked at about 1.6x. The proteosome inhibitor MG132 blocked this dequenching, indicating that dequenching was largely due to proteosomes rather than lysosomes. As we did for eGFP synthesis, we also multiplied the rate data by the extract concentration to infer a degradation rate for endogenous proteins, where both the substrate and the proteolysis machinery would be affected by cytoplasmic concentration; this shifted the activity peak to ∼1.8x (Figure 3D). Note that by paired *t*-test, the average for the 1.8x data was not significantly higher than the average for the 2x data (*p* = 0.26 for a one-tailed *t*-test), so the optimal cytoplasmic concentration for DQ-BSA dequenching may actually be higher than 1.8x. The specific activity calculation (Figure 3E) showed that the enzyme activity fell above ∼1.4x cytoplasmic concentration. At lower cytoplasmic concentrations, the relationship between this gauge of activity and concentration was complicated; perhaps simple bimolecular kinetics do not pertain in this regime.

**Figure 3.**
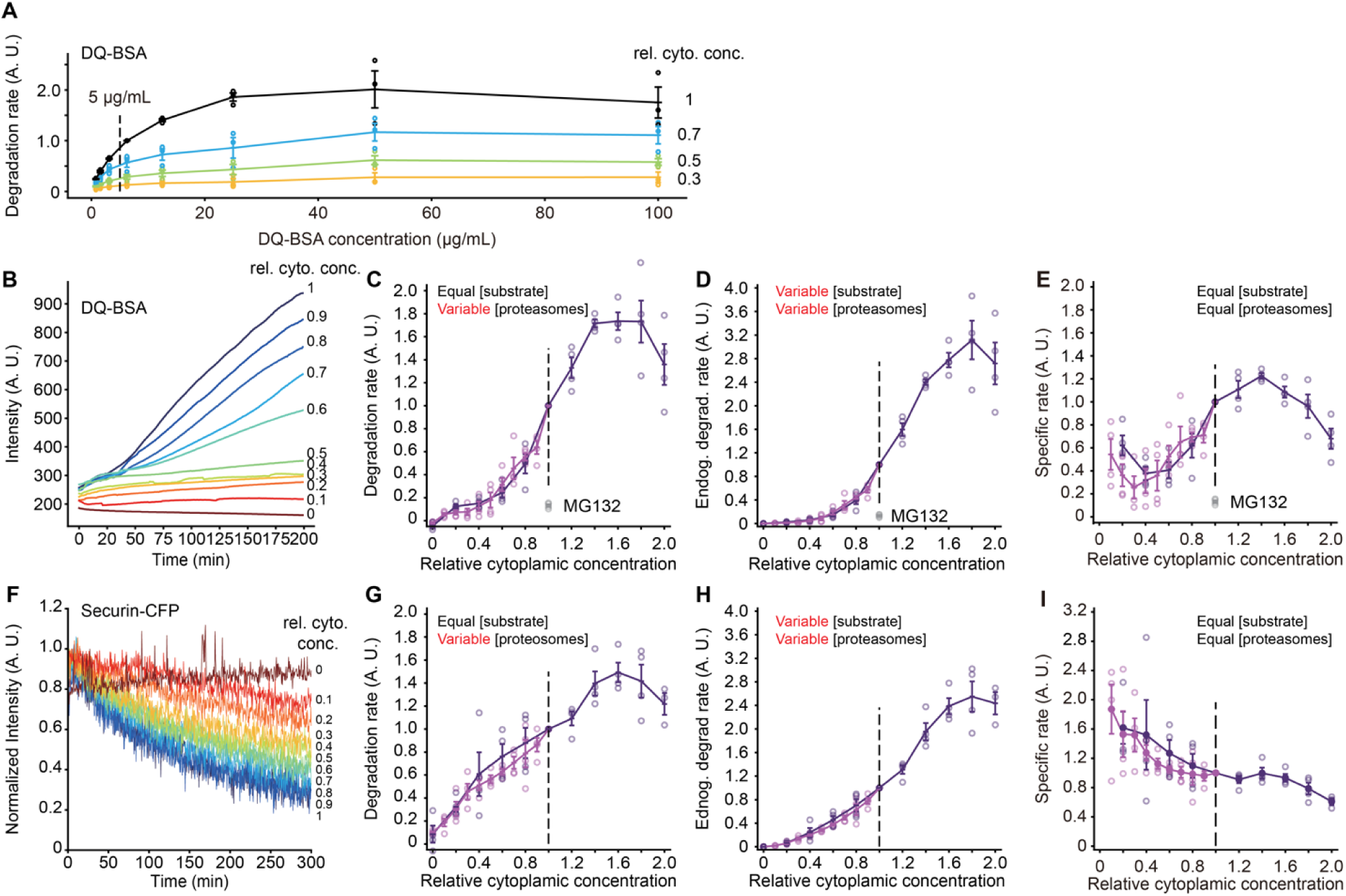
The rate of protein degradation is maximal at cytoplasmic concentrations of ∼1.8x. (A) Titration of substrate protein concentration for DQ-BSA degradation experiments. The indicated concentration (5 µg/mL) was chosen for the experiments in (B-E). (B) DQ-BSA fluorescence as a function of time for various dilutions of a 1x extract. (C) Degradation rate as a function of cytoplasmic concentration. These are the directly-measured data from experiments where the DQ-BSA concentration was kept constant and the proteolysis machinery was proportional to the cytoplasmic concentration. The grey data points denoted MG132 are from 1x extracts treated with 200 µM MG132, a proteosome inhibitor. Data are from 4 experiments for dilution from 1x extracts and 4 experiments for dilution from 2x retentates. Data are normalized relative to the degradation rates at a cytoplasmic concentration of 1x. Means and standard errors are overlaid on the individual data points. In this and the subsequent panels, the darker purple represents data from diluting 2x retentates and the lighter purple from diluting 1x extract. (D) Inferred degradation rates for the situation where the substrate concentration as well as the proteosome concentration is proportional to the cytoplasmic concentration. The rates from (C) were multiplied by the relative cytoplasmic concentrations. (E) Inferred degradation rates for the situation where both the substrate concentration and the proteosome concentration are kept constant at all dilutions. The rates from (C) were divided by the relative cytoplasmic concentrations. (F) Degradation of securin-CFP as a function of time for various dilutions of a 1x extractt. (G) Degradation rate as a function of cytoplasmic concentration. These are the directly-measured data from experiments where the securin-CFP concentration was kept constant but the proteosome concentration was proportional to the cytoplasmic concentration. Data are from 4 experiments for dilution from 1x extracts and 4 experiments for dilution from 2x retentates. Data are normalized relative to the degradation rates at a cytoplasmic concentration of 1x. Means and standard errors are overlaid on the individual data points. (H) Inferred degradation rate for the situation where both the substrate and proteosome concentrations are proportional to the cytoplasmic concentration. The rates from (G) were multiplied by the relative cytoplasmic concentrations. (I) Inferred degradation rates for the situation where both the substrate and the proteosome concentration are kept constant at all dilutions. The rates from (G) were divided by the relative cytoplasmic concentrations.

As a second measure of protein degradation, we used a securin-CFP fusion protein as a reporter. Securin is a cell cycle protein known to be targeted for proteasome-mediated protein degradation by the anaphase promoting complex/cyclosome (APC/C) in late mitosis. We measured the decay of securin-CFP fluorescent intensity and calculated the degradation rate (Kamenz et al., 2021) for each of the dilution conditions (Figure 3F). As was the case with DQ-BSA dequenching, the protein degradation rate peaked at a ∼1.6x cytoplasmic macromolecular concentration (Figure 3G), and the inferred rate for a substrate being diluted along with the degradation machinery peaked at 1.8x (Figure 3H). Again, by paired *t*-test, the average for the 1.8x data was not significantly higher than the average for the 2x data (*p* = 0.34 for a one-tailed *t*-test), so the optimal cytoplasmic concentration for securin-CFP degradation may be higher than 1.8x. The inferred specific activity for securin-CFP degradation fell steadily with concentration (Figure 3I). Thus, by both measures, protein degradation rates were maximal at a cytoplasmic concentration of about 1.8x, higher than the optimal concentration for protein synthesis.

### Viscosity affects protein translation rate and, to a lesser extent, protein degradation

The decrease in translation at high cytoplasmic macromolecule concentrations suggests that translation is diffusion-controlled in a viscous cytoplasm, and that concentrated cytoplasm has a higher viscosity that reduces the molecular movement necessary for the translation reaction. To test these ideas, we first measured how diffusion coefficients vary with cytoplasmic concentration. We used single particle tracking of fluorescently labeled PEGylated 100 nm diameter polystyrene beads, which are comparable in size to some of the large complexes involved in translation (Figure 4A). In 1x extracts, the motion of the beads was subdiffusive, with the diffusivity exponent α to be 0.88 ± 0.06 (means ± S.E.) (Figure 4B, C). We calculated an average effective diffusion coefficient *D* from a fit of the random walk diffusion equation to the data over a time scale of 1 s. This was found to be 0.36 ± 0.23 μm^2^/s (means ± S.E.), in good agreement with previous studies (Delarue et al., 2018; Huang et al., 2022). There was substantial variability from position to position in the speed of diffusion (Figure 4B, insets, and 4C).

**Figure 4.**
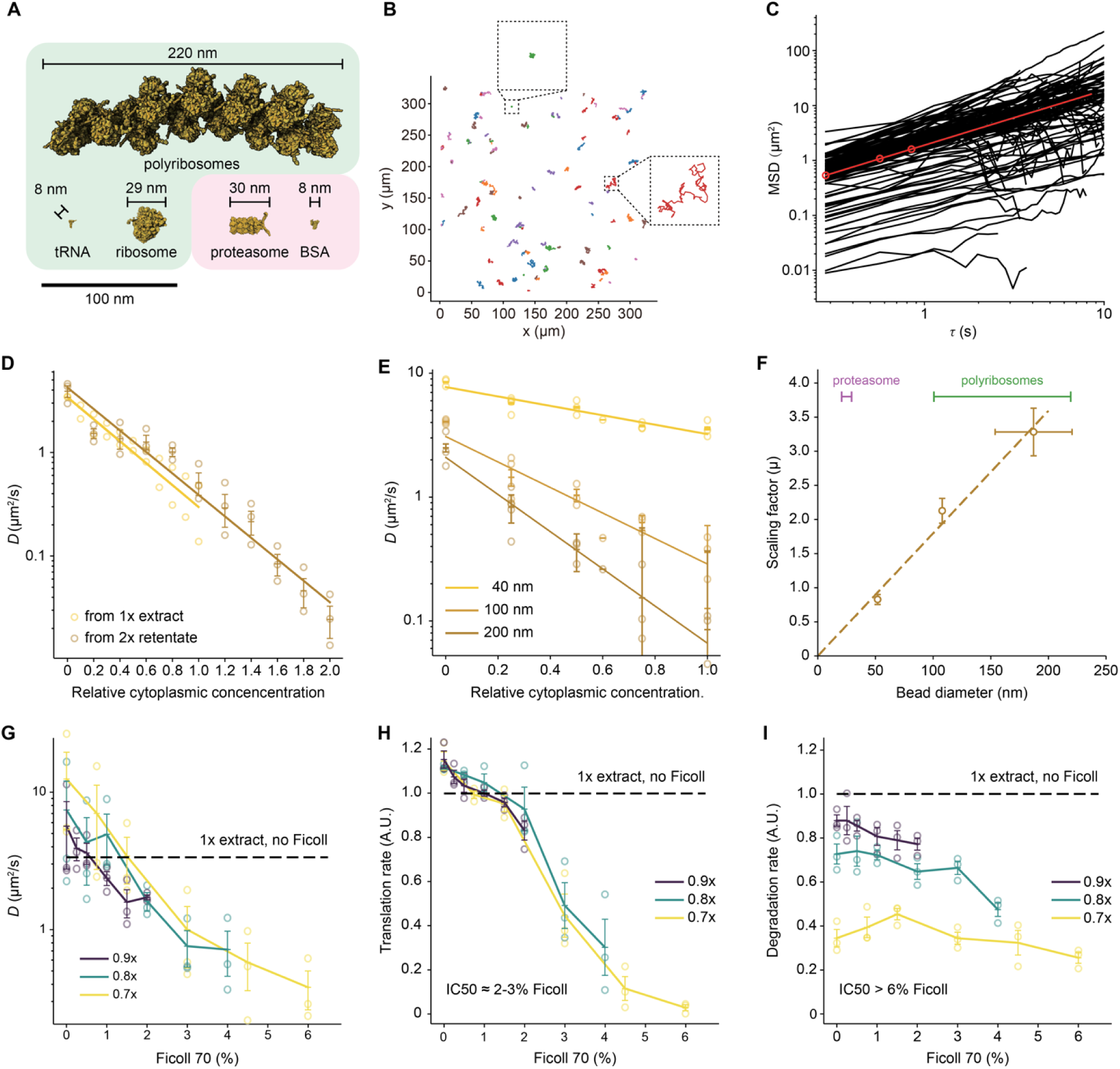
The effect of cytoplasmic concentration on diffusion, and the effect of Ficoll 70 on translation and protein degradation. (A) The sizes of various macromolecules and complexes involved in translation and degradation. (B) Single particle traces for diffusion of 100 nm fluorescent beads in 1x cytoplasmic extracts. Two examples of location-to-location variability are highlighted. (C) Mean-squared displacement for 110 individual trajectories (black) and average mean-squared displacement (red) as a function of the time difference τ. Effective diffusion coefficients were calculated from the first 1 s of data. (D) Effective diffusion coefficients for 100 nm fluorescent beads as function of relative cytoplasmic concentration. Data are from 3 experiments for the 2x extract dilution and from 2 experiments for the 1x extract dilution. Error bars for the 2x extract dilution represent means ± standards errors. (E) Effective diffusion coefficients for beads of different diameter (nominally 40 nm, 100 nm, and 200 nm) as a function of relative cytoplasmic concentration. Data are from 3 experiments. Means and standard errors are overlaid on the individual data points. (F) The scaling factor µ (from Eq. 1) as a function of bead diameter. The apparent bead diameters (nominally 40, 100, and 200 nm) were calculated from their diffusion coefficients in extract buffer with no sucrose using the Stokes-Einstein relationship. Scaling factors are from 3 experiments and are shown as means ± S.E. Bead diameters are from 3 experiments for the 40 nm beads and 4 experiments for the 100 and 200 nm beads, and again are plotted as means ± S.E. The diameters of proteosomes and polyribosomes are shown for comparison. (G) Diffusion coefficients of 40 nm beads as a function of Ficoll 70 concentration. Extracts were prepared at 0.7x, 0.8x, and 0.9x as indicated and supplemented with Ficoll to yield the final concentrations (w/vol) shown on the *x*-axis. Data are from 3 experiments. Means and standard errors are overlaid on the individual data points. Diffusion coefficients for the undiluted 1x extracts were also measured and the average is shown for reference. (H) Translation rates, using the eGFP assay, as a function of Ficoll 70 concentration. Extracts were prepared at 0.7x, 0.8x, and 0.9x as indicated and supplemented with Ficoll to yield the final concentrations (w/vol) shown on the *x*-axis. Data are from the same 3 experiments shown in (G). Means and standard errors are overlaid on the individual data points. Translation rates for the undiluted 1x extracts were also measured and the average is shown for reference. (I) Degradation rates, using the DQ-BSA assay, as a function of Ficoll 70 concentration. Extracts were prepared at 0.7x, 0.8x, and 0.9x as indicated and supplemented with Ficoll to yield the final concentrations (w/vol) shown on the *x*-axis. Data are from the same 3 experiments shown in (G). Means and standard errors are overlaid on the individual data points. Degradation rates for the undiluted 1x extracts were also measured and the average is shown for reference.

Similarly high variability has been reported for the diffusion of genetically-encoded nanoparticles expressed in *S. pombe* (Garner et al., 2023), which is thought to reflect the heterogeneity of the cytoplasmic environment.

The effective diffusion coefficient for the 100 nm beads varied dramatically with cytoplasmic concentration; it was ∼11x higher in filtrate (0x cytoplasm) compared to 1x extracts, and ∼11x lower in 2x extracts. The measured diffusion coefficients obeyed Phillies’s law (Phillies, 1986; Phillies, 1988):

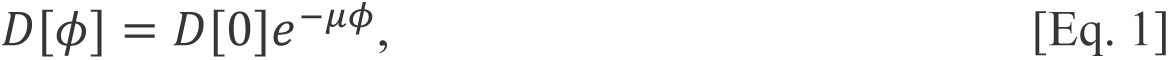

where *#x1D437;[#x1D719;]* is the diffusion coefficient at a relative cytoplasmic concentration *#x1D719;*, *#x1D437;[0]* is the diffusion coefficient in filtrate, and *µ* is a scaling factor that depends upon the size of the probe. A similar relationship between diffusion coefficients and macromolecular concentration has been observed for large multimeric protein complexes (Alric et al., 2022; Molines et al., 2022).

Similarly, the scaling factor varied approximately linearly with the measured size of the beads (Figure 4D) (Alric et al., 2022). Thus, diffusion was markedly affected by changing the cytoplasmic macromolecule concentration over a 0x to 2x range, and the changes were greatest for large probe particles.

Next, we diluted extracts to 0.7x, 0.8x, or 0.9x, and altered the cytoplasmic viscosity by adding Ficoll 70, a protein-sized (70 kDa) carbohydrate that can act both as a crowding agent and a viscogen (Figure 4G). The effective diffusion coefficients were found to decrease with increasing Ficoll concentration; 6% Ficoll 70 decreased the diffusion coefficient for a 40 nm bead by a factor of 33, and 2-3% Ficoll 70 yielded diffusion coefficients comparable to those measured in 2x cytoplasm. Thus, Ficoll can be used to bring about the marked changes in viscosity seen when cytoplasm is concentrated, without changing protein concentration.

We therefore asked whether this range of Ficoll 70 concentrations would inhibit protein synthesis and degradation. Note that Ficoll might be expected to have either of two opposite effects on enzyme reaction rates: by acting as a crowding agent, it increases the concentrations of the reactants and thus could increase the rate of a reaction; but by acting as a viscogen, it could slow protein motions and decrease in the reaction rate. In vitro studies have shown that either of these effects can predominate (Aoki et al., 2013; Minton, 2001; Tan et al., 2013). We found that the rate of translation of eGFP monotonically decreased with increasing Ficoll 70 concentration, with an IC50 of ∼2-3% (Figure 4H). Diffusion coefficients in these Ficoll-supplemented 0.7x to 0.9x extracts supplemented with 2-3% Ficoll 70 were similar to those seen in 2x extracts with no Ficoll. Thus protein synthesis is sensitive to viscosity over a range relevant to the cytoplasmic concentration/dilution experiments.

The rate of DQ-BSA unquenching was substantially less sensitive to viscosity (Figure 4I), with an IC50 greater than 6% Ficoll 70. Thus, both protein synthesis and protein degradation, as measured by the eGFP and DQ-BSA assays, are sensitive to the viscosity of the cytoplasm, with synthesis being more sensitive than degradation. This difference in sensitivity is sufficient to account for the different optimal cytoplasmic concentrations found for translation and degradation.

### A model for the effect of diffusion on reaction rates

Final, we asked whether we could derive a simple model to account for the observed rates of protein translation and degradation as a function of cytosolic protein concentration. In particular, we wished to see if plausible assumptions could account for the biphasic nature of the curves, and to explore how differences in the kinetics of translation vs. degradation might account for the differences in their concentration optima.

We assume that the rate determining reaction for each process is a mass action bimolecular reaction:

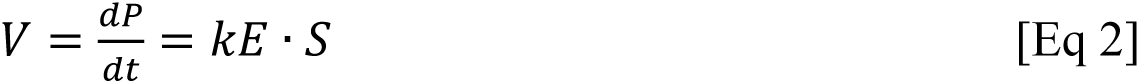

where *V* is the rate of the reaction, *P* is the product of the reaction, *E* is the enzyme, *S* the substrate, and *P* the product of the reaction. Alternatively, we could assume a two-step, saturable mechanism (see Supplementary Information), but the mass action treatment suffices for present purposes and is simpler. The enzyme and substrate concentrations are linearly proportional to the relative cytoplasmic concentration *#x1D719;*. We can therefore write:

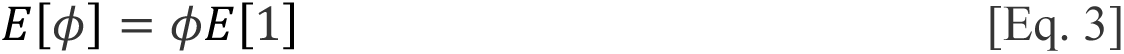

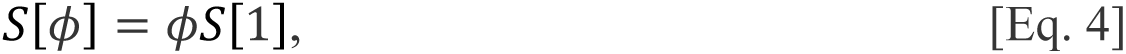

where *#x1D438;[#x1D719;]* and *#x1D446;[#x1D719;]* are the enzyme and substrate concentrations at a relative cytoplasmic concentration of *#x1D719;*, and *#x1D438;[1]* and *#x1D446;[1]* are the enzyme and substrate concentrations at a relative cytoplasmic concentration 1x. Substituting into Eq. 2 yields:

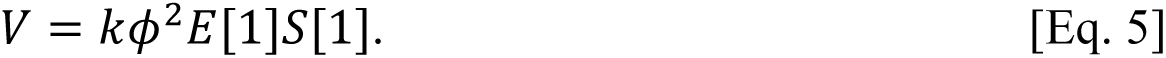

Next we consider the dependence of the rate constant on the cytoplasmic protein concentration. From the Smoluchowski equation (Smoluchowski, 1917), the collision rate and the association rate constant *#x1D458;* are proportional to the sum of the diffusion coefficients of *E* and *S*, and from Phillies’s law (Eq. 1) (Phillies, 1986; Phillies, 1988), we take the diffusion coefficients to be negative exponential functions of the cytoplasmic protein concentration:

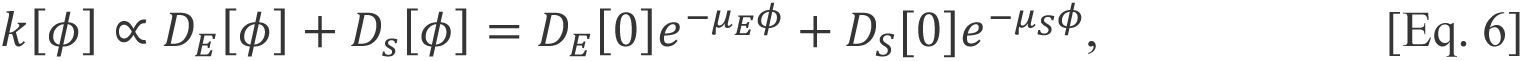

where *#x1D437;_E_[0]* and *#x1D437;_S_[0]* are the diffusion coefficients of the enzyme and the substrate at a cytoplasmic concentration of 0 (i.e. in filtrate), and the *µ*’s are scaling factors. For the special case where either one diffusion coefficient is much smaller than the other, or the scale factors are equal, we can simplify this to:

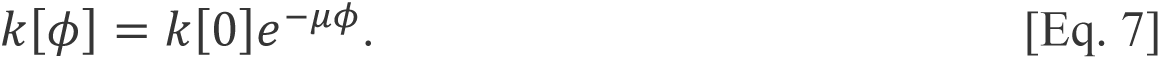

Note that it is a bit awkward to consider a value of *#x1D458;* at a relative cytoplasmic concentration of 0, since the reacting species are cytoplasmic macromolecules that are absent from the filtrate. We can avoid this by instead writing:

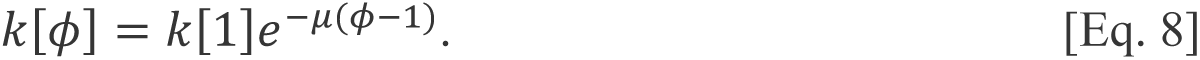

It follows that:

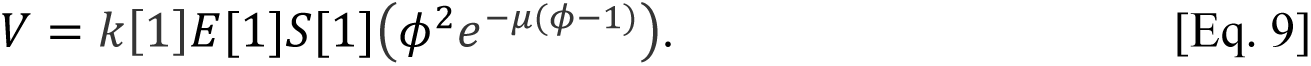

Finally, we can make use of the fact that the scaling factor *µ* is linearly proportional to the size of the diffusing particle *d_p_*, and write an expression that explicitly acknowledges particle size:

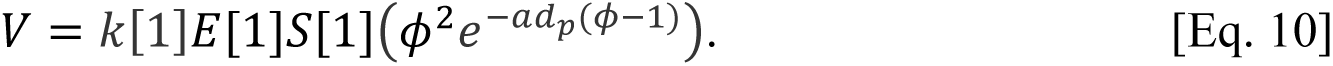

where *a* is a new scaling factor that relates *d_p_* to *µ*. Eq. 10 describes how the rate of a mass action, bimolecular reaction whose reactants obey Phillies’s law and mass would be expected to vary with the cytoplasmic macromolecule concentration and molecular diameter. If we measure rates in arbitrary units with the rate at *#x1D719; = 1* taken as 1, then:

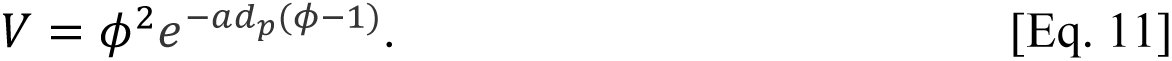

Note that we have an experimental estimate for *a* (which, from Figure 4F is 0.018 nm^-1^), leaving only one adjustable parameter, *d_p_*, the macromolecular diameter. Eq. 11 defines a biphasic, non monotonic curve (Figure 5A), and the larger the assumed macromolecular diameter, the further to the left the curve’s maximum lies (Figure 5A). For a given value of *d_p_*, the optimal cytoplasmic concentration is given by:

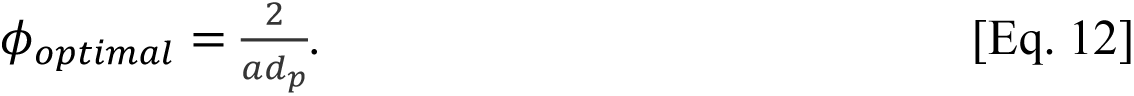

**Figure 5.**
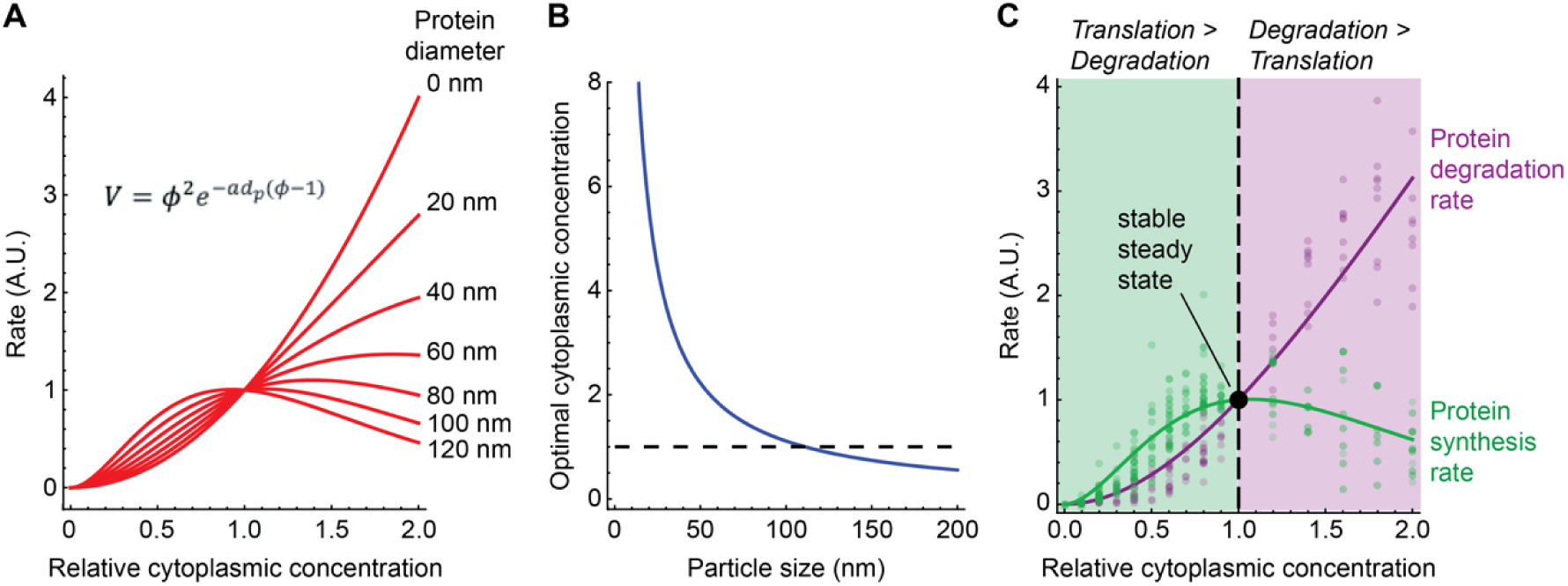
Homeostasis in a model of the effect of cytoplasmic concentration of translation and protein degradation. (A) Plot of Eq. 11, which relates a bimolecular reaction rate to the relative cytoplasmic concentration, for various sizes of proteins. We assumed *a* = 0.018 nm^-1^ (from Figure 4F). (B) Calculated optimal relative cytoplasmic concentration for proteins of different assumed sizes, again assuming *a* = 0.018 nm^-1^ (C) Fits of Eq. 11 to the experimental data for translation (green) and degradation (purple) as a function of cytoplasmic concentration, calculated assuming that both the substrate and enzyme varied with the cytoplasmic concentration. All of the data from Figures 2D, H and 3D, H were included in the fits. The *#x1D445;^2^* values are 0.92 for the translation data and 0.95 for the degradation data. The fitted values for the size of the proteins involved are 104 ± 2 nm (mean ± S.E.) for translation and 14 ± 1 nm (mean ± S.E) for degradation. The fitted optimal cytoplasmic concentrations are 1.07 ± 0.02 for translation and 8.1 ± 0.8 for degradation (mean ± S.E.).

The experimentally observed rates for translation and degradation are well captured by Eq. 11, with the fitted values of *d_p_* being 104 ± 2 nm for translation and 14 ± 1 nm for degradation (means ± S.E.) (Figure 5C).

The result is a dynamical system that will be in steady state—the translation and degradation rates will be equal—at a relative cytoplasmic concentration of 1x. Moreover, the steady state is guaranteed to be stable. If the system is perturbed such that the cytoplasmic concentration exceeds 1x, then the degradation rate will rise and the translation rate will fall, driving the system back toward the 1x steady state (Figure 5C). Conversely, if the cytoplasmic concentration falls below 1x, the translation rate will exceed the synthesis rate, again driving the system back toward the physiological set point (Figure 5C).

## Discussion

Here we have tested the hypothesis that the concentration of macromolecules in the cytoplasm is set to maximize the rates of important biochemical reactions. We found that in cycling *Xenopus* egg extracts, the rate of translation, as measured by the synthesis of eGFP from an exogenous mRNA and the incorporation of ^35^S-methionine into endogenous translation products, does peak at a 1x cytoplasmic concentration, consistent with the maximal speed hypothesis (Figure 2). This finding fits well with previous studies of cost minimization and near-optimal resource allocations in models of *E. coli* protein synthesis (Hu et al., 2020; Klumpp et al., 2013; Scott et al., 2014).

However, protein degradation, as measured by DQ-BSA dequenching and CFP-securin degradation peaks at a higher cytoplasmic concentration, ∼1.8x (Figure 3). The difference in concentration optima can be explained by the differences in the sensitivities of translation vs. degradation to viscosity: the viscogen Ficoll 70 is more potent in inhibiting translation than degradation (Figure 4H, I). This in turn may be due to differences in the sizes of the macromolecular complexes involved in the two processes, as the diffusion coefficients for large fluorescent beads are more affected by cytoplasmic concentration than those of smaller beads (Figure 4E, F). Other factors that might make translation be more diffusion-limited than protein degradation could also contribute the differential sensitivity of the two processes to changes in cytoplasmic concentration. The different optimal cytoplasmic concentrations for translation vs. degradation mean that the system is homeostatic: increasing the concentration of macromolecules in the cytoplasm would increase the rate of degradation and decreases the rate of translation, whereas decreasing the cytoplasmic concentration would decrease the rate of degradation (Figure 5). This behavior can be explained through a theoretical treatment based on mass action kinetics and Phillies’s law (Phillies, 1986; Phillies, 1988).

Both the mRNA translation and protein degradation rates measured here were found to smoothly vary with cytoplasmic concentration (Figures 2 and 3). Thus, there was no evidence for critical concentrations above or below which the processes abruptly ceased. We suspect that any such critical points lie outside the range of concentrations examined here. Likewise, the effects of high cytoplasmic concentration on translation and degradation dynamics appeared to be largely reversible, since the activities measured in a 2x extract diluted back to 1x were very similar to those in the original 1x extract (Figures 2 and 3).

Although the conventional wisdom is that almost all biochemical reactions are reaction controlled (i.e., the catalytic rate constant *k*_2_ is much smaller than *k*_-1_) rather than diffusion controlled (*k*_-1_ is much smaller than *k*_2_) (Bar-Even et al., 2011; Davidi et al., 2016), both of the processes measured here were inhibited by the crowding agent/viscogen Ficoll 70, with translation substantially more sensitive than degradation. The simplest interpretation is that translation (in particular) is running relatively close to the diffusion limit, possibly because in the extract, the diffusion of large macromolecules required for dissociation (*k*_-1_) is quite slow.

Note that the shapes of the rate curves mean that protein synthesis and degradation can be viewed as a negative feedback system. Increasing the cytoplasmic protein concentration increases the cytoplasmic viscosity, which negatively affects translation, which makes the protein concentration eventually drop—a negative feedback loop based on the sensitivity of translation to viscosity. Conversely, decreasing the cytoplasmic protein concentration decreases the cytoplasmic viscosity, which mitigates the drop in translation rates that would normal be expected from a decrease in ribosome and mRNA concentration. Such feedback regulation may also pertain to the oscillation of biomass growth rate and maintenance of cytoplasmic density found in single mammalian cells (Liu et al., 2020). Negative feedback is a common theme in homeostatic systems. This process can alternatively be viewed as a variation on end product inhibition, where the product of protein synthesis inhibits translation, but through the intermediacy of changes in protein diffusion rates rather than through the direct binding of the product to the enzyme.

We do not yet have direct evidence for the time scale of this proposed protein concentration homeostasis. Based on the measurement of 43 h for the median half-life of a *Xenopus* protein during embryogenesis (Peshkin et al., 2015), we suspect that the response might require tens of hours. This time scale would be particularly appropriate for protein homeostasis in the immature oocyte, the egg’s immediate precursor in development. The oocyte is thought to live for weeks or months in the frog ovary and to vary little in terms of size, appearance, and composition during this time (Ferrell, 1999; Smith et al., 1991). In contrast, the changes in protein synthesis and degradation are likely to be too slow make a significant impact during the decrease in cytoplasmic concentration that occurs during mitosis, since embryonic mitosis is only about 15 min in duration.

## Acknowledgements

This work was supported by a grant from the NIH (R35 GM131792) to J.E.F.

## Author contributions

Conceptualization, Y.C., J.H., and J.E.F.; Methodology, Y.C., J.H, C.P., and J.E.F.; Software, Y.C.; Formal Analysis, Y.C. and J.E.F.; Investigation, Y.C.; Resources, C.P.; Writing – Original Draft, Y.C. and J.E.F.; Visualization, Y.C. and J.E.F.; Supervision, J.E.F.; Funding Acquisition, J.E.F.

## Star Methods

### Extract preparation

Cycling extracts were prepared as described previously (Chang and Ferrell, 2018; Murray, 1991) with the following modifications. Briefly, freshly laid frog eggs were collected, washed with 20 g/L L-cysteine pH 7.8, and incubated for 3-5 min to remove the jelly coat. The eggs were then washed twice with ∼150 mL 0.2x 6s (MMR) solution (20 mM NaCl, 1 mM HEPES, 400 µM KCl, 400 µM CaCl2, 200 µM MgCl2, and 20 µM EDTA pH 7.8) and resuspended in 50 mL 0.2x MMR solution. Calcium ionophore A23187 (C7522, Sigma) was added to a final concentration of 0.5 µg/mL to activate the eggs. After 2 min of activation, liquid was removed, and the eggs were washed twice with ∼150 mL 0.2xMMR solution and three times with ∼150 mL extract buffer [100 mM KCl, 50 mM sucrose, 10 mM HEPES pH 7.7 (with KOH), 1 mM MgCl2, and 100 µM CaCl2]. Twenty min after activation (>80% of the eggs showed contraction of the animal pole), the eggs were transferred to a 14 mL round-bottom polypropylene tube (352059, Corning) and packed for 1 min at 300 × *g*. Excess liquid on top of the eggs was removed, and the egg-containing tube was chilled on ice. The eggs were crushed by centrifugation at 16,000 × *g* for 15 min at 4°C. The cytoplasmic layer was then collected by puncturing the side of the extract-containing tube at ∼2 mm above the interphase between the extract layer and the yolk layer. The extract was allowed to flow into a collecting Eppendorf tube by gravity or by gently pressing the tube opening with one finger to create a positive pressure inside the tube. The collected extract was mixed with 10 µg/mL leupeptin, 10 µg/mL pepstatin, 10 µg/mL chymostatin, and 10 µg/mL cytochalasin B, and further refined by centrifugation at 16,000 × *g* for 5 min at 4°C using a tabletop refrigerated centrifuge. After the refining centrifugation, the clarified extract was transferred to a new tube.

CSF extracts (Kamenz et al., 2021; Murray, 1991) were prepared similarly to cycling extracts, with the differences being that the eggs were washed with ∼150 mL CSF extract buffer (100 mM KCl, 50 mM sucrose, 10 mM potassium HEPES pH 7.7, 5 mM EGTA pH 7.7, 2 mM MgCl2, and 0.1 mM CaCl2) four times immediately after dejellied, without 0.2x MMR washes or calcium ionophore activation. The whole process between dejellying and the crushing spin typically took ∼10 min. After the refining centrifugation, the extract was transferred to a new tube and supplemented with 100 µg/mL cycloheximide.

### Filtrate and retentate preparation

Extract (400 μL) was transferred to a 10 kDa molecular weight cut-off centrifugal filter unit (UFC501096, Millipore) placed in a collection tube and centrifuged at 16,000 × *g* for 10 min at 4°C three times. After each centrifugation period, the extract was taken out of the centrifuge and mixed by pipetting. The filtrate was collected in the collection tube, and the retentate was collected from the filtration unit.

### SiR-tubulin intensity

To monitor microtubules in cycling extracts, 200 nM SiR-tubulin was added to the extract, retentate, and filtrate. Then the extract or retentate was mixed with different volume fractions of filtrate to create a series of dilution conditions. Five µL of the dilutions were loaded onto a 96 well polystyrene assay plate (3368, Corning) and gently spread using the pointed end of the loading pipette tip to allow even coverage of extract at the bottom of the well. A layer of 100 µL heavy mineral oil (330760, Sigma) was pipetted to cover the extract and prevent evaporation.

The 96 well plate was immediately loaded onto an inverted epi-fluorescence microscope (DMI8, Leica) for imaging at a frame rate of 0.5 min^-1^. The median intensity from the center 1/9 of each frame was measured and plotted with off-sets in Figures 1D and 1E.

### eGFP translation and DQ-BSA degradation

For eGFP translation and DQ-BSA degradation experiments, the extract, filtrate, or retentate was mixed with 2.5 ng/µL (unless otherwise stated) eGFP mRNA (L-7201-100, Trilink Biotechnologies) or 5 ng/µL DQ-BSA (D12050, Thermo Fisher) on ice. The extract or retentate was mixed with different proportions of the filtrate to generate different dilutions. The dilutions (15 µL) were then added to a clear bottom 384-well plate (324021, Southern Labware) and equilibrated to room temperature. The imaging plate was then loaded onto an inverted fluorescence microscope for time course measurements at a frame rate of 1 min^-1^ or 0.5 min^-1^.

To calculate the rate of eGFP protein synthesis and DQ-BSA degradation, the median intensity of the center quarter of each frame was measured. The raw rates were extracted by calculating the slope of a linear segment from the intensity-time plot. The linear segment was typically between 50 and 120 min for eGFP translation, and between 25 and 120 min for DQ-BSA degradation experiments. Intensity trajectories were manually inspected and the linear segments were adjusted individually to ensure linearity. The raw rates from dilutions of 1x extract (or 2x retentate) were normalized by the rate measured in the original 1x extract (or the reconstituted 1x extract) from the same batch of eggs to control for variability due to batch variation of the eggs and experimental conditions, allowing comparison among experiments.

### ^35^S-methionine labeling

A cycling extract was concentrated as described above. The extract, retentate, and filtrate were supplemented with 1% v/v ^35^S-methionine to a final concentration of ∼0.5 µCi/µL. The filtrate was then mixed with different volume fractions of extract or retentate to generate a series of dilutions. For the “CHX” sample, 100 µg/mL cycloheximide was added to a 1x extract. The extract was then sampled at 15-min intervals, and the translation process was halted by directly mixing 5 µL samples with 100 µL H2O, which was then mixed with 100 µL 50% TCA (trichloroacitic acid, T0699, Sigma). To collect and clean up the TCA precipitable material, 50 µL of the homogeneous extract/TCA mixture was passed through a glass fiber filter (WHA1820025, Sigma) prewetted with 5% TCA, then 1 mL of 5% TCA was passed through the filter to remove soluble material, and the filter was washed with 2 mL of 95% ethanol and dried on vacuum. The filter with collected material was dropped into a 20-mL scintillation vial (03- 337-2, Thermo Fisher) containing 10 mL scintillation fluid (111195, RPI Research Products).

The radioactivity was measured using a liquid scintillation counter. The rate was calculated similarly to the eGFP translation and DQ-BSA degradation experiments.

### Securin-CFP degradation

The securin-CFP degradation experiments followed a previous protocol (Kamenz et al., 2021) with modifications. The fluorescent probe, securin-CFP, was made by mixing 10 µg of an SP6- securin-CFP plasmid in 20 µL H2O with 30 µL SP6 High-Yield Wheat Germ Protein Expression System (TnT® L3261, Promega), and incubating at room temperature for 2 h per the manufacturer’s instructions. CSF extracts were used for these experiments. The CSF extract was additionally supplemented with purified recombinant nondegradable Δ90 sea urchin cyclin B protein at a concentration capable of driving the extract into an M-phase arrest and incubated at room temperature for 30 min. 0.8 mM CaCl2 was added to the extract and incubated for an additional 30 min to degrade endogenous cyclin B. The extract was then divided into two fractions, one kept on ice and the other concentrated using the previously stated method. A series of dilutions were reconstituted by mixing the filtrate with either the extract kept on ice or retentate from the concentrator. 19 µL of the extracts were added and mixed with 1 µL of the in vitro transcribed and translated securin-CFP and pipette into a glass-bottomed 384-well plate. As a background control, we also included a well of extract with no securin-CFP. The time courses of fluorescence intensity were recorded using a fluorescence plate reader at a rate of 2 min^-1^.

To calculate the degradation rate, each experimental reading was subtracted by the corresponding background measurement. The first few data points typically increased with time, possibly due to equilibration of the fluorophore. Therefore, instead of normalizing to the first data point in each time series, the background corrected intensities were divided by the maximum of the first 15 time points (7.5 min). The normalized intensities were fitted to *A* = *A*[0] e^-*kt*^ +C, where *A* is the fluorescence, *k* is the rate constant, and *t* is time, and *A*[0] (constrained to be greater than 0.95), *k*, and C (constrained between 0 and 0.05) are fitting variables. The value of *k* measured for each dilution condition from 1x extract (or from 2x retentate) was normalized to 1x extract (or nominal 1x reconstituted from the 2x retentate) from the same experiment to control for variations among batches of eggs.

### Single particle tracking

PEGylated fluorescent particles were prepared by mixing 50 µL of 20 mg/mL methoxypolyethylene glycol amine 750 (07964, Sigma), 5 µL of fluorescent polystyrene nano beads (2% solid, F8888 and F8795, Thermo Fisher), 50 µL of 30 mM N-hydroxysulfosuccinimide (56485, Sigma) in 200 mM borate buffer pH8.2, and 10 µL of 100 mM N-(3-dimethylaminopropyl)-N′-ethylcarbodiimide hydrochloride (03450, Sigma). The PEGylation reaction was incubated at room temperature for 20 h. The reaction was then stopped by 1:100 dilution with water, dialysis with 3 M NaCl, and three subsequent dialyses with water. The particles were diluted further to make it so that a 1:100 dilution provided a suitable concentration for segmentation and tracking. 1:100 (v:v) of the beads were mixed into the extract by pipetting. 5 µL of the extract was placed in the center of a well in a glass-bottom 96-well plate. The plate was loaded onto an inverted epifluorescence microscope. The extract was allowed to equilibrate with the environment for 5 min, and a movie was taken at the appropriate wavelength at a frame rate of 3 Hz using a 40x objective.

The time-lapse videos of particle movements were analyzed with a custom Python script. Briefly, the images were flat field corrected (background flat fields were generated using BaSiC, an ImageJ package) and bleach corrected. The particles were called and linked using the Trackpy library with adaptive mode, which allows calling particle movements with large step size variations. Typical starting parameters for Trackpy were: diameter = 15, maxsize = 7, minmass = 650, search_range =30, ecc_threshold = 1, percentile = 99.5, topn = 300, memory = 1; drift correction was typically off unless the movie had a translational flow. Movies were discarded if they contained a strong convergent or divergent flow. Parameters were adjusted for individual movies to allow capturing the greatest number of tracks without sacrificing tracking quality. The mean squared displacement for an individual trajectory was calculated by 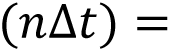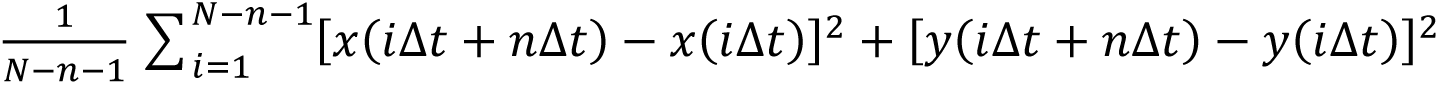, where *N* is the number of frames in a trajectory and *x* and *y* are the coordinates at *#x1D456;Δ#x1D461;* or *#x1D456;Δ#x1D461; + #x1D45B;Δ#x1D461;*. The ensemble mean squared displacement was calculated by*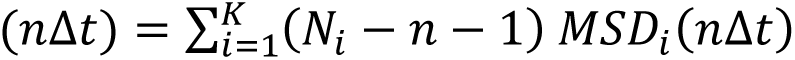*, where *K* is the *i=1* number of trajectories, *Ni* is the number of frames for the *i*^th^ trajectory, and *MSDi*(*nΔt*) is the individual MSD of the *i*^th^ trajectory for a time lag of *nΔt*.

### Estimation of the effective diffusion coefficient for the time scale of 1 s

To calculate the effective diffusion coefficient, *Deff*, a linear fit was made to the first 3 values to the ensemble MSD vs τ plot (corresponding to τ = 1/3, 2/3, and 1s). The slope of the fitted line was calculated to obtain MSD/τ. The effective diffusion coefficient was calculated by *Deff* = MSD/(4 τ).

### Particle size estimation

PEGylated particles were resuspended in an extract buffer without sucrose. The effective diffusion coefficient for each type of particle was measured by particle tracking as above. The diameters of the particles were calculated using a rearrangement of the Stokes-Einstein equation: *dpar* = *kB T*/ (3*πηDeff*), where *kB* is the Boltzmann constant (1.380649ꞏ10^-23^ N m K^-1^), *T* is temperature (296.15 K), and *η* is the viscosity of extract buffer without sucrose (assumed to be similar to water at 0.001 N m^-2^ s).

## Supplementary Information

**Figure S1.**
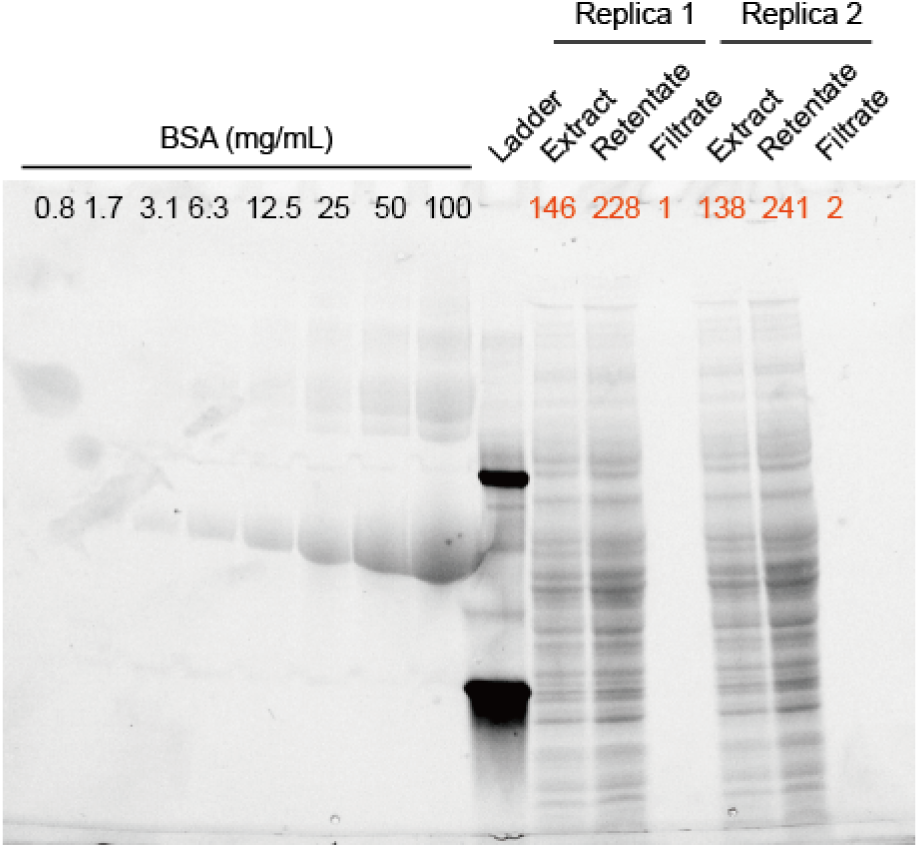
Trihalo compound-stained PAGE gel of proteins from 1x extract, filtrate, and 2x retentate A representative trihalo compound-stained PAGE gel showing a BSA standard and two biological replicates of extract, retentate, and filtrate. The estimated protein concentrations in mg/mL (using the BSA standard as a reference) for the extract, retentate, and filtrate samples are shown in orange. However, it should be noted that the trihalo compound used in staining the gel depends on tryptophan residues to fluoresce. Since BSA has only 0.3% tryptophan residues (compared to 1% in proteins overall), the protein concentrations in the extract, retentate, and filtrate samples are likely overestimated. By Bradford assay the typical protein concentration of extracts was 50-70 mg/mL.

**Figure S2.**
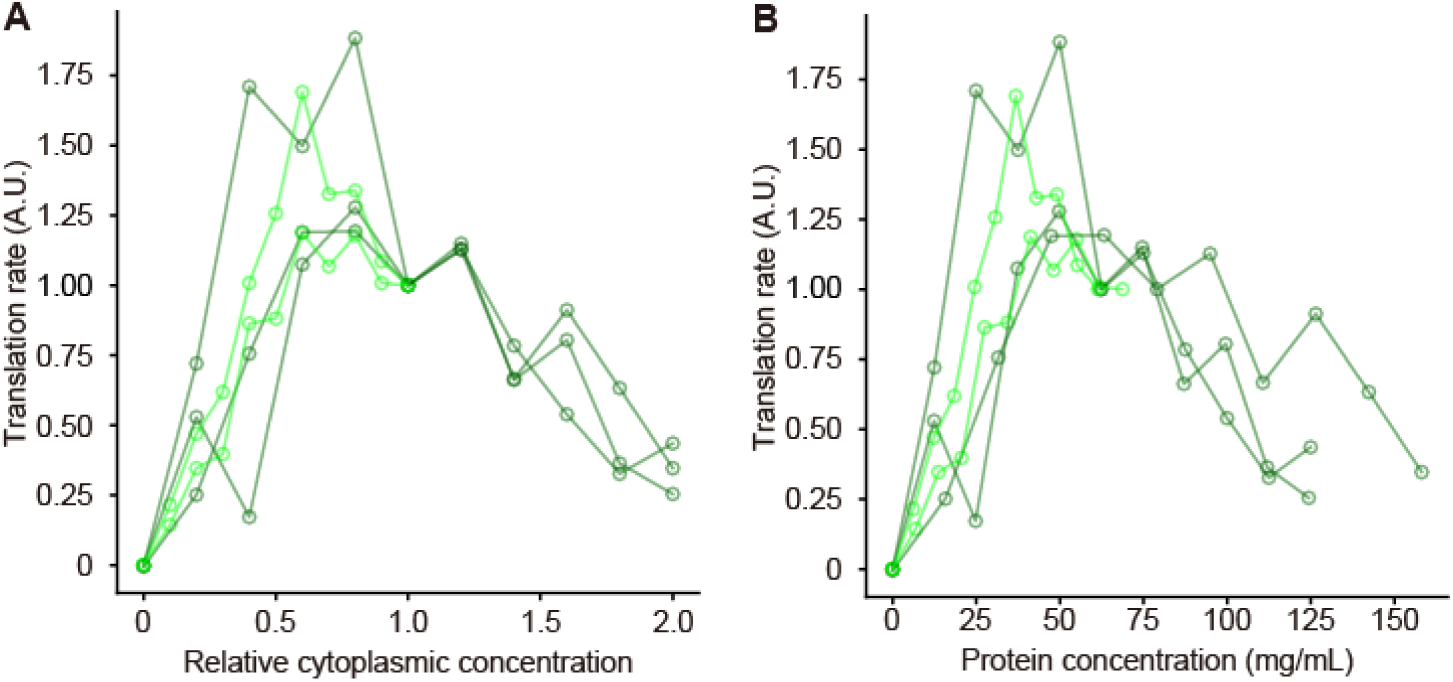
Using nominal vs. measured protein concentrations to compare translation rates in different experiments. (A) Translation rate as a function of nominal cytoplasmic concentration. These are the directly measured data from experiments where the eGFP mRNA concentration was kept constant and the translation machinery was proportional to the cytoplasmic concentration. Data points from same experiment are connected. Data are normalized relative to the translation rates at a cytoplasmic concentration of 1x. Relative cytoplasmic concentration are assumed to be 1.0 for 1x extract and 2.0 for the 2x retentate. (B) Translation rate as a function of measured protein concentration. These are the directly measured data from experiments where the eGFP mRNA concentration was kept constant and the translation machinery was proportional to the cytoplasmic concentration. Data points from same experiment are connected. Data are normalized relative to the translation rates at a cytoplasmic concentration of 1x. Protein concentrations were measured for the starting extract and the retentate, instead of assuming that they were 1x and 2x. Protein concentrations for the dilutions were calculated from these the respective starting material. Note that the experiment-to experiment variation is similar regardless of whether nominal or measured protein concentrations are used.

**Figure S3.**
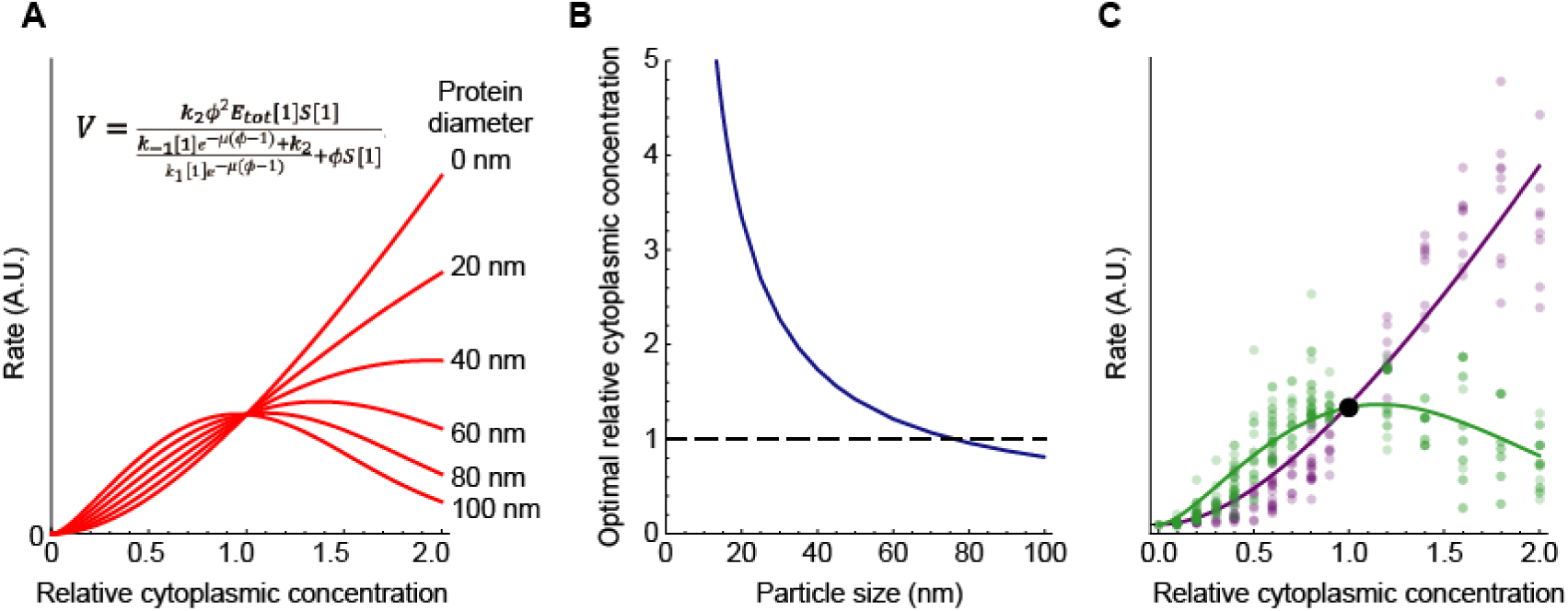
Homeostasis in a Michaelis-Menten model of the effect of cytoplasmic concentration of translation and protein degradation. (A) Plot of Eq. S10, which relates a bimolecular reaction rate to the relative cytoplasmic concentration, for various sizes of proteins, assuming that Michaelis-Menten kinetics are relevant. We assumed *a* = 0.018 nm^-1^ (from Figure 4F) and arbitrarily chose the following values for the parameters: *#x1D438;_tot_[1] = #x1D446;[1] = #x1D458;_1_[1] = #x1D458;_-1_[1] = #x1D458;_2_[1] = 1*. (B) Calculated optimal relative cytoplasmic concentration for proteins of different assumed sizes, assuming the same parameters. (C) Fits of Eq. S10 to the experimental data.

### Movie S1. Effects of cytoplasmic concentration on self-organization and cycling in dilutions from 1x extract

SiR-tubulin staining (left) and SiR-tubulin fluorescence intensity as a function of time (right) in an extract after various dilutions. The same data are shown in Figure 1D. The starting material was a 1x extract, diluted with various proportions of filtrate and imaged in a 96-well plate under mineral oil. All fields are shown at equal exposure. The fluorescence intensities shown on the right were quantified from the center 1/9 of the wells and subsequently rescaled using the maximum and minimum values to enhance the variability in the intensities, particularly in cases of low cytoplasmic concentration.

### Movie S2. Effects of cytoplasmic concentration on self-organization and cycling in dilutions from 2x retentate

SiR-tubulin staining (left) and SiR-tubulin fluorescence intensity as a function of time (right) in an extract after various dilutions. The data were collected as in Movie S1 except that the starting material was a 2x retentate. The same data are shown in Figure 1E.

### Movie S3. Cell-like compartment formation and cell cycle oscillation in 0.2x and 0.3x cytoplasm

SiR-tubulin staining was conducted on a 0.2x extract (left) and a 0.3x extract (right) as shown in Movie S1 and Figure 1D. The fluorescence intensities were rescaled to improve the visibility of the SiR-tubulin staining, and are displayed at a higher magnification,

### A model for the effect of diffusion on reaction rates assuming Michaelis-Menten kinetics

In the main text, we derived a simple model to account for the observed rates of protein translation and degradation as a function of cytosolic protein concentration assuming a mass action kinetics. Here, we assume a Michaelis-Menten enzymatic reaction kinetics to generate response curves, and test if such an assumption also accounts the biphasic nature of the response curves of protein translation and degradation vs cytoplasmic concentration.

To start, we assume that the rate determining reaction for each process is a biomolecular enzymatic reaction:

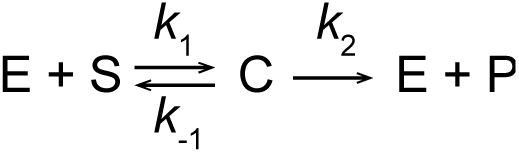

where E is the enzyme, S the substrate, C the enzyme-substrate complex, and P the product of the reaction. If we assume that the system is in steady state, with 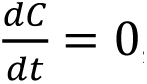, and that the substrate concentration is much higher than the enzyme concentration, then the rate of this process is described by the Michaelis-Menten equation:

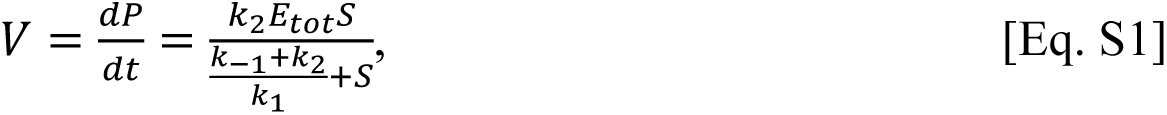

where #x1D438;_tot_ = #x1D438;+ #x1D436;.

Next we want to add cytoplasmic concentration dependence to the terms on the right-hand side of Eq. 2. The enzyme and substrate concentrations are linearly proportional to the relative cytoplasmic concentration *#x1D719;*. We can therefore write:

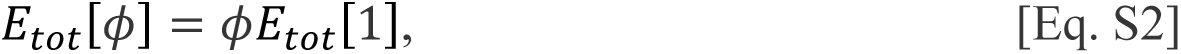

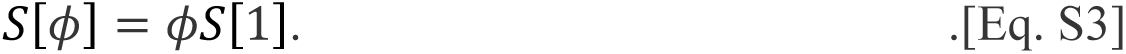

Substituting into Eq. 2 yields:

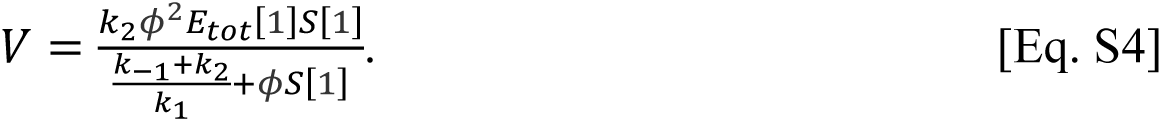

Again, from the Smoluchowski equation (Smoluchowski, 1917) and Phillies’s law (Eq. 1) (Phillies, 1986; Phillies, 1988), we take the diffusion coefficients to be negative exponential functions of the cytoplasmic protein concentration:

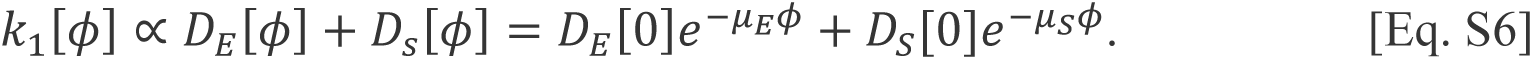

For the special case where either one diffusion coefficient is much smaller than the other, or the scale factors are equal, we can simplify this to:

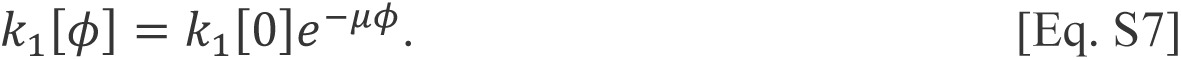

Similarly, we can rewrite Eq. S7 as:

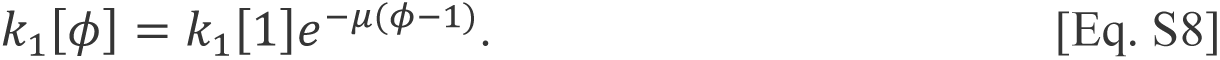

We assume that *#x1D458;_-1_*varies with the diffusion coefficients and the cytoplasmic concentration in the same way, and that *#x1D458;_2_*, the rate constant for the catalytic step, is independent of the cytoplasmic concentration. It follows that:

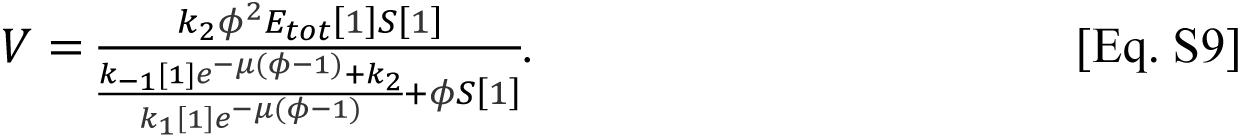

Finally, since the scaling factor *µ* is linearly proportional to the size of the diffusing particle *dp*, we can write an expression that explicitly acknowledges particle size:

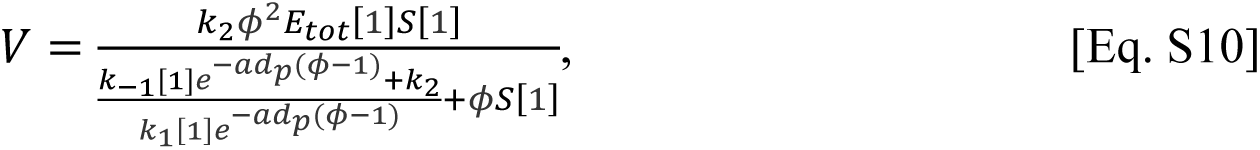

where *a* is a new scaling factor that relates *dp* to *µ*. Eq. S10 describes how the rate of a Michaelis-Menten reaction whose reactants obey Phillies’s law and the Smoluchowski equation would be expected to vary with the cytoplasmic macromolecule concentration and molecular diameter.

We do not have experimental estimates for most of the parameters in Eq. S10 (except for *a*, which, from Figure 4F is 0.018 nm^-1^) for either translation or protein degradation. However, we can get a feel for Eq. S10 by arbitrarily assuming some parameter values (*#x1D438;_tot_[1] = #x1D446;[1] = #x1D458;_1_[1] = #x1D458;_-1_[1] = #x1D458;_2_[1] = 1*) and plotting *V* as a function of *#x1D719;* for macromolecules of different assumed macromolecular diameters. The equation defines a biphasic, non-monotonic curve (Figure S3A), and the larger the assumed macromolecular radius, the further to the left the curve’s maximum lies (Figure S3B). The observed rates for translation and degradation are fairly well captured by assuming that the relevant macromolecules are 78 nm for translation and 0 nm for degradation (Figure S3C).

## References

1. Phillies, G.D. (1986). Universal scaling equation for self-diffusion by macromolecules in solution. Macromolecules 19, 2367–2376.

2. Phillies, G.D.J. (1988). Quantitative prediction of *α* in the scaling law for self-diffusion. Macromolecules 21, 3101–3106.

3. Smoluchowski, M. (1917). Mathematical theory of the kinetics of the coagulation of colloidal solutions. Zeitschrift für Physikalische Chemie 92, 129–168.

## References

1. Afanzar, O., Buss, G.K., Stearns, T., and Ferrell, J.E., Jr. (2020). The nucleus serves as the pacemaker for the cell cycle. Elife 9, e59989.

2. Alric, B., Formosa-Dague, C., Dague, E., Holt, L.J., and Delarue, M. (2022). Macromolecular crowding limits growth under pressure. Nature Physics 18, 411–416.

3. Amodeo, A.A., Jukam, D., Straight, A.F., and Skotheim, J.M. (2015). Histone titration against the genome sets the DNA-to-cytoplasm threshold for the *Xenopus* midblastula transition. Proc Natl Acad Sci U S A 112, E1086–1095.

4. Aoki, K., Takahashi, K., Kaizu, K., and Matsuda, M. (2013). A quantitative model of ERK MAP kinase phosphorylation in crowded media. Sci Rep 3, 1541.

5. Bar-Even, A., Noor, E., Savir, Y., Liebermeister, W., Davidi, D., Tawfik, D.S., and Milo, R. (2011). The moderately efficient enzyme: evolutionary and physicochemical trends shaping enzyme parameters. Biochemistry 50, 4402–4410.

6. Chang, J.B., and Ferrell, J.E., Jr. (2013). Mitotic trigger waves and the spatial coordination of the *Xenopus* cell cycle. Nature 500, 603–607.

7. Chang, J.B., and Ferrell, J.E., Jr. (2018). Robustly Cycling *Xenopus laevis* Cell-Free Extracts in Teflon Chambers. Cold Spring Harb Protoc 2018, pdb.prot097212.

8. Cheng, X., and Ferrell, J.E., Jr. (2019). Spontaneous emergence of cell-like organization in *Xenopus* egg extracts. Science 366, 631–637.

9. Davidi, D., Noor, E., Liebermeister, W., Bar-Even, A., Flamholz, A., Tummler, K., Barenholz, U., Goldenfeld, M., Shlomi, T., and Milo, R. (2016). Global characterization of in vivo enzyme catalytic rates and their correspondence to in vitro *kcat* measurements. Proc Natl Acad Sci U S A 113, 3401–3406.

10. Delarue, M., Brittingham, G.P., Pfeffer, S., Surovtsev, I.V., Pinglay, S., Kennedy, K.J., Schaffer, M., Gutierrez, J.I., Sang, D., Poterewicz, G., et al. (2018). mTORC1 Controls Phase Separation and the Biophysical Properties of the Cytoplasm by Tuning Crowding. Cell 174, 338–349 e320.

11. Dill, K.A., Ghosh, K., and Schmit, J.D. (2011). Physical limits of cells and proteomes. Proc Natl Acad Sci U S A 108, 17876–17882.

12. Drenckhahn, D., and Pollard, T.D. (1986). Elongation of actin filaments is a diffusion limited reaction at the barbed end and is accelerated by inert macromolecules. Journal of Biological Chemistry 261, 12754–12758.

13. Dunphy, W.G., Brizuela, L., Beach, D., and Newport, J. (1988). The *Xenopus* cdc2 protein is a component of MPF, a cytoplasmic regulator of mitosis. Cell 54, 423–431.

14. Elowitz, M.B., Surette, M.G., Wolf, P.E., Stock, J.B., and Leibler, S. (1999). Protein mobility in the cytoplasm of *Escherichia coli*. Journal of bacteriology 181, 197–203.

15. Fang, F., and Newport, J.W. (1991). Evidence that the G1-S and G2-M transitions are controlled by different cdc2 proteins in higher eukaryotes. Cell 66, 731–742.

16. Ferrell, J.E., Jr. (1999). *Xenopus* oocyte maturation: new lessons from a good egg. Bioessays 21, 833–842.

17. Fox, C.A., Sheets, M.D., and Wickens, M.P. (1989). Poly(A) addition during maturation of frog oocytes: distinct nuclear and cytoplasmic activities and regulation by the sequence UUUUUAU. Genes Dev 3, 2151–2162.

18. Garner, R.M., Molines, A.T., Theriot, J.A., and Chang, F. (2023). Vast heterogeneity in cytoplasmic diffusion rates revealed by nanorheology and Doppelganger simulations. Biophys J 122, 767–783.

19. Gires, P.-Y., Thampi, M., Krauss, S.W., and Weiss, M. (2023). Exploring generic principles of compartmentalization in a developmental in vitro model. Development 150.

20. Green, S.L. (2009). The laboratory Xenopus sp, 1 edn (CRC Press).

21. Hu, X.P., Dourado, H., Schubert, P., and Lercher, M.J. (2020). The protein translation machinery is expressed for maximal efficiency in Escherichia coli. Nat Commun 11, 5260.

22. Huang, W.Y.C., Cheng, X., and Ferrell, J.E., Jr. (2022). Cytoplasmic organization promotes protein diffusion in *Xenopus* extracts. Nat Commun 13, 5599.

23. Jin, M., Tavella, F., Wang, S., and Yang, Q. (2022). In vitro cell cycle oscillations exhibit a robust and hysteretic response to changes in cytoplasmic density. Proc Natl Acad Sci U S A 119, e2109547119.

24. Kamenz, J., Qiao, R., Yang, Q., and Ferrell, J.E., Jr. (2021). Real-Time Monitoring of APCAnaphase-promoting complex (APC)/C-Mediated Substrate Degradation Using *Xenopus laevis* Egg Extracts. In Cell Cycle Oscillators : Methods and Protocols, A.S. Coutts, and L. Weston, eds. (New York, NY: Springer US), pp. 29–38.

25. Kanki, J.P., and Newport, J.W. (1991). The cell cycle dependence of protein synthesis during *Xenopus laevis* development. Dev Biol 146, 198–213.

26. King, R.W., Deshaies, R.J., Peters, J.M., and Kirschner, M.W. (1996). How proteolysis drives the cell cycle. Science 274, 1652–1659.

27. Klumpp, S., Scott, M., Pedersen, S., and Hwa, T. (2013). Molecular crowding limits translation and cell growth. Proc Natl Acad Sci U S A 110, 16754–16759.

28. Liu, X., Oh, S., and Kirschner, M.W. (2022). The uniformity and stability of cellular mass density in mammalian cell culture. Front Cell Dev Biol 10, 1017499.

29. Liu, X., Oh, S., Peshkin, L., and Kirschner, M.W. (2020). Computationally enhanced quantitative phase microscopy reveals autonomous oscillations in mammalian cell growth. Proc Natl Acad Sci U S A 117, 27388–27399.

30. Meneau, F., Dupre, A., Jessus, C., and Daldello, E.M. (2020). Translational Control of *Xenopus* Oocyte Meiosis: Toward the Genomic Era. Cells 9, 1502.

31. Milo, R., Jorgensen, P., Moran, U., Weber, G., and Springer, M. (2009). BioNumbers— the database of key numbers in molecular and cell biology. Nucleic Acids Research 38, D750–D753.

32. Minshull, J., Blow, J.J., and Hunt, T. (1989). Translation of cyclin mRNA is necessary for extracts of activated *Xenopus* eggs to enter mitosis. Cell 56, 947–956.

33. Minton, A.P. (2001). The influence of macromolecular crowding and macromolecular confinement on biochemical reactions in physiological media. The Journal of biological chemistry 276, 10577–10580.

34. Mitchison, T.J., and Field, C.M. (2021). Self-Organization of Cellular Units. Annu Rev Cell Dev Biol 37, 23–41.

35. Molines, A.T., Lemiere, J., Gazzola, M., Steinmark, I.E., Edrington, C.H., Hsu, C.T., Real-Calderon, P., Suhling, K., Goshima, G., Holt, L.J., et al. (2022). Physical properties of the cytoplasm modulate the rates of microtubule polymerization and depolymerization. Dev Cell 57, 466–479 e466.

36. Murray, A., W. (1991). Cell Cycle Extracts. In Methods in Cell Biology, Brian K. Kay, and H.B. Peng, eds. (Academic Press), pp. 581–605.

37. Murray, A.W., and Kirschner, M.W. (1989). Cyclin synthesis drives the early embryonic cell cycle. Nature 339, 275–280.

38. Neurohr, G.E., and Amon, A. (2020). Relevance and Regulation of Cell Density. Trends in cell biology 30, 213–225.

39. Newport, J., and Kirschner, M. (1982a). A major developmental transition in early *Xenopus* embryos: I. characterization and timing of cellular changes at the midblastula stage. Cell 30, 675–686.

40. Newport, J., and Kirschner, M. (1982b). A major developmental transition in early *Xenopus* embryos: II. Control of the onset of transcription. Cell 30, 687–696.

41. Newport, J., and Spann, T. (1987). Disassembly of the nucleus in mitotic extracts: membrane vesicularization, lamin disassembly, and chromosome condensation are independent processes. Cell 48, 219–230.

42. Oh, S., Lee, C., Yang, W., Li, A., Mukherjee, A., Basan, M., Ran, C., Yin, W., Tabin, C.J., Fu, D., et al. (2022). Protein and lipid mass concentration measurement in tissues by stimulated Raman scattering microscopy. Proc Natl Acad Sci U S A 119, e2117938119.

43. Paris, J., Osborne, H.B., Couturier, A., Le Guellec, R., and Philippe, M. (1988). Changes in the polyadenylation of specific stable RNA during the early development of *Xenopus laevis*. Gene 72, 169–176.

44. Peshkin, L., Wuhr, M., Pearl, E., Haas, W., Freeman, R.M., Jr., Gerhart, J.C., Klein, A.M., Horb, M., Gygi, S.P., and Kirschner, M.W. (2015). On the Relationship of Protein and mRNA Dynamics in Vertebrate Embryonic Development. Dev Cell 35, 383–394.

45. Phillies, G.D. (1986). Universal scaling equation for self-diffusion by macromolecules in solution. Macromolecules 19, 2367–2376.

46. Phillies, G.D.J. (1988). Quantitative prediction of *α* in the scaling law for self-diffusion. Macromolecules 21, 3101–3106.

47. Richter, K., Nessling, M., and Lichter, P. (2007). Experimental evidence for the influence of molecular crowding on nuclear architecture. Journal of cell science 120, 1673–1680.

48. Rosenthal, E.T., Tansey, T.R., and Ruderman, J.V. (1983). Sequence-specific adenylations and deadenylations accompany changes in the translation of maternal messenger RNA after fertilization of Spisula oocytes. J Mol Biol 166, 309–327.

49. Ruderman, J.V., Woodland, H.R., and Sturgess, E.A. (1979). Modulations of histone messenger RNA during the early development of *Xenopus laevis*. Dev Biol 71, 71–82.

50. Schwanhausser, B., Busse, D., Li, N., Dittmar, G., Schuchhardt, J., Wolf, J., Chen, W., and Selbach, M. (2011). Global quantification of mammalian gene expression control. Nature 473, 337–342.

51. Scott, M., Klumpp, S., Mateescu, E.M., and Hwa, T. (2014). Emergence of robust growth laws from optimal regulation of ribosome synthesis. Mol Syst Biol 10, 747.

52. Smith, L.D., Xu, W.L., and Varnold, R.L. (1991). Oogenesis and oocyte isolation. Methods Cell Biol 36, 45–60.

53. Smoluchowski, M. (1917). Mathematical theory of the kinetics of the coagulation of colloidal solutions. Zeitschrift für Physikalische Chemie 92, 129–168.

54. Son, S., Kang, J.H., Oh, S., Kirschner, M.W., Mitchison, T.J., and Manalis, S. (2015). Resonant microchannel volume and mass measurements show that suspended cells swell during mitosis. Journal of Cell Biology 211, 757–763.

55. Tan, C., Saurabh, S., Bruchez, M.P., Schwartz, R., and Leduc, P. (2013). Molecular crowding shapes gene expression in synthetic cellular nanosystems. Nat Nanotechnol 8, 602–608.

56. Zimmerman, S.B., and Trach, S.O. (1991). Estimation of macromolecule concentrations and excluded volume effects for the cytoplasm of *Escherichia coli*. J Mol Biol 222, 599–620.

57. Zlotek-Zlotkiewicz, E., Monnier, S., Cappello, G., Le Berre, M., and Piel, M. (2015). Optical volume and mass measurements show that mammalian cells swell during mitosis. Journal of Cell Biology 211, 765–774.

